# The misplaced mouse Pax6 interneuron subclass: A cross-species transcriptomic reassignment

**DOI:** 10.1101/2025.07.23.666342

**Authors:** Jonathan M Werner, Hamsini Suresh, Jesse Gillis

## Abstract

Subclass cell-types are strongly conserved across mammalian brains, making the systemic absence of a caudal ganglionic eminence (CGE) Pax6 interneuron subclass in mouse transcriptional atlases unexpected. Through cross-species transcriptomic analysis, we identify a Pax6 subclass homolog in mouse and uncover primate-specific divergence in the Sncg subclass. Our results highlight the pitfalls of relying on single marker genes and provide evolution-aware annotations and marker sets to support robust cross-mammal interneuron comparisons.

## Main

Identifying homologous neural cell types across species is central to studying brain evolution^1^. In single-cell transcriptomics, this is complicated by technical variability and species-specific shifts in gene expression that hinder accurate cross-species cell-type matching^2–4^. Despite this, subclass-level cell-types consistently emerge as robust and replicable across platforms and species^5–10^, establishing them as a reliable framework for comparative neuroscience.

A notable exception is the CGE Pax6 interneuron subclass. While consistently annotated in human^8,11–13^ and non-human primate atlases^7,14^, it is conspicuously absent from mouse datasets^9,15–18^, including comprehensive whole-brain atlases^19,20^ (Fig. 1A). In mouse, PAX6 marker-expression is inconsistently associated with clusters labeled as CGE Lamp5, Vip, or Sncg subclasses^15–19^. This raises a central question: is the Pax6 subclass a primate-specific innovation, or has its murine homolog been misclassified?

**Figure 1:**
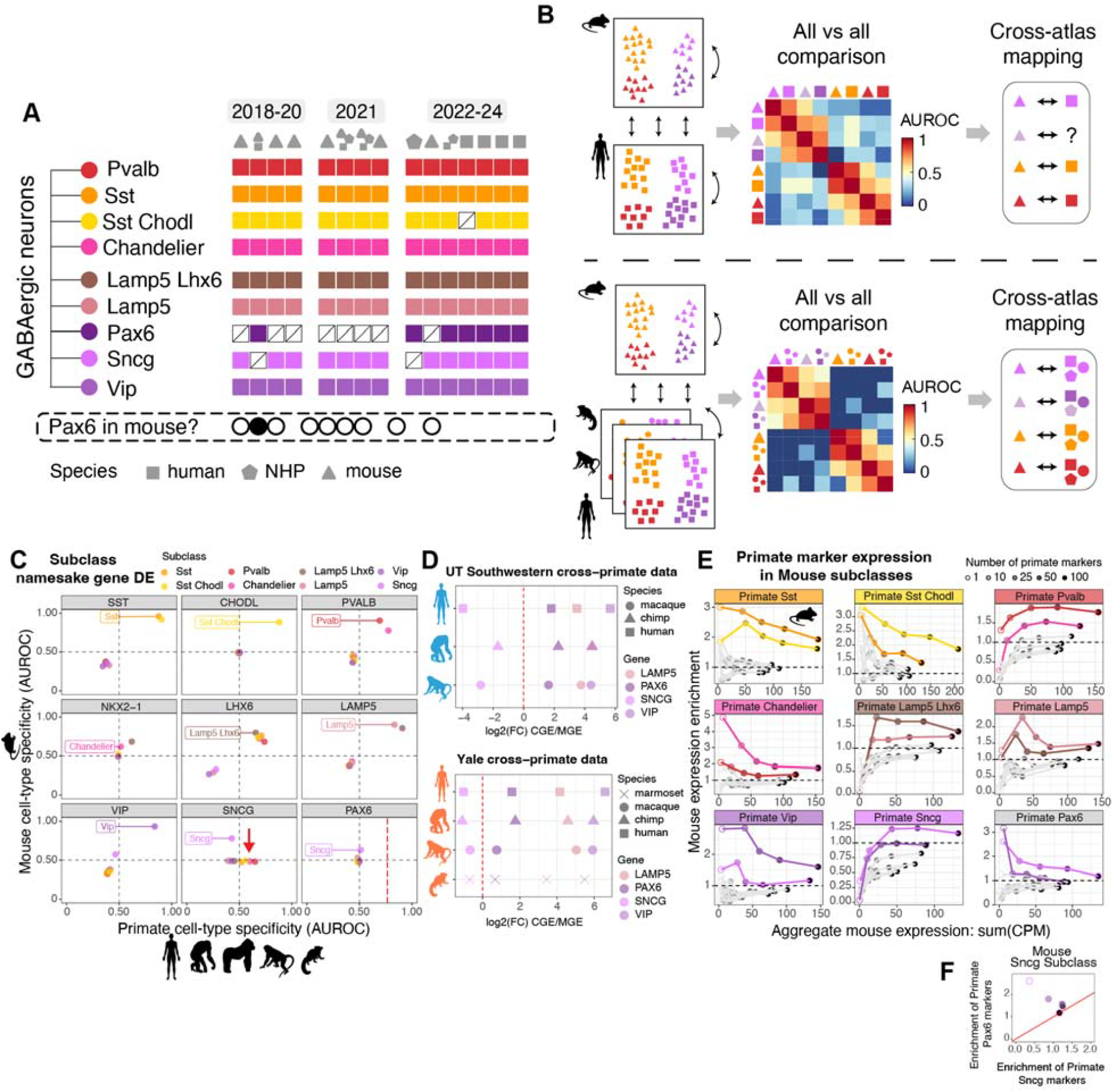
Primate-mouse subclass differential expression comparisons suggest a Pax6-subclass misannotation rather than absence in mouse datasets. **A:** Schematic depicting subclass-level annotations for mouse, non-human primate (NHP) and human single cell datasets. Pax6 subclass annotations are consistently absent in mouse datasets, yet are consistently present in NHP and human datasets. **B:** Schematic depicting how cross-species consensus cell-type annotations can help resolve cell-type alignment ambiguities when performing pair-wise species comparisons. **C:** Scatter plots comparing primate (Allen MTG dataset) and mouse differential expression (DE, MetaMarker AUROCs) for the namesake genes of the interneuron subclass annotations. The red arrow is highlighting the MGE-positive DE of the SNCG gene in primates. The red dotted line is depicting the primate PAX6 AUROC in the primate Pax6 subclass, due to the absence of Pax6-subclass annotations in mouse. **D:** Dot plots comparing the CGE/MGE log2 fold-change of the LAMP5, PAX6, SNCG, and VIP genes across the primate species for the UT Southwestern and Yale datasets. Positive CGE DE is to the right of the dotted line. **E:** Scatter-line plots comparing the aggregate mouse expression (summation of CPM values) against the mouse expression enrichment (see *Cell-type marker and MetaMarker generation* methods section) for increasing numbers of top primate subclass MetaMarkers (Supp. Table 1). Only the top 2 mouse subclasses per primate subclass are shown in color, determined by their maximal expression of the primate markers. **F:** Scatter plot comparing the expression enrichment of the primate Pax6 and Sncg subclass markers for the mouse Sncg subclass.

To address this, we performed a meta-analysis of CGE and MGE interneuron subclasses across diverse mouse and primate datasets (see methods). Our data collection includes mouse and primate (human, chimpanzee, gorilla, macaque, marmoset) interneurons from both 10x and Smart-seq platforms, single-cell and single-nucleus preparations, and 18 mouse and 3 primate brain regions. Data were drawn from the Allen Institute (mouse and primate), Broad Institute (mouse), Yale (primate), and UT Southwestern (primate), enabling a broad evolutionary comparison to resolve cell-type annotation ambiguities (Fig. 1B).

We started mouse and primate subclass comparisons by leveraging previously established cross-primate consensus interneuron subclasses from 5 primate species^6,7^ (Allen Institute data, Supp. Figs. 1-2). This prior work identified nine conserved primate interneuron subclasses: Pvalb, Chandelier, Sst, Sst Chodl, Lamp5, Lamp5 Lhx6, Vip, Sncg, and Pax6. We then curated a high-confidence subset of mouse data by first identifying replicable cell clusters across our 8 diverse Allen and Broad Institute mouse datasets using MetaNeighbor^21^, which utilizes gene co-expression similarity to identify replicable cell-types across datasets (Supp. Fig. 3, see methods).

We then compared differential expression (DE) of namesake marker genes for each subclass, including PAX6, to assess whether Pax6-like cells are absent or misannotated in mouse (Fig. 1C). Several markers show expected conserved DE across species (PVALB, SST, CHODL, LAMP5, LHX6, VIP), but we also identify striking species-specific patterns for the Chandelier^22^ and Sncg subclass namesake genes, though most dramatically for SNCG (Fig. 1C). SNCG is a strong marker for the mouse Sncg subclass, but is unexpectedly positively DE in primate MGE-derived rather than CGE-derived subclasses. A strong transcriptomic separation of CGE- and MGE-derived subclasses is a consistent finding across studies^5–8,11,15–18^, marking the apparent MGE switch in SNCG DE within primates as particularly surprising. We confirm the MGE-positive DE of SNCG in both the Yale and UT Southwestern primate datasets (Fig. 1D, Supp. Fig. 4), validating a significant shift in DE patterns between mouse and primates. Importantly, we also observe mild positive DE of PAX6 in the mouse Sncg subclass, suggesting misannotation rather than a true loss of Pax6-like interneurons in mouse.

The CGE-to-MGE inversion of SNCG underscores how even well-established subclass markers can diverge across species, challenging the reliability of single or small marker sets for cross-species annotation. To test the robustness of subclass identity, we quantified the enrichment and aggregate expression of increasing numbers of primate subclass markers (Supp. Table 1) in mouse data. While top individual markers can show weak or inconsistent differential expression in the expected mouse subclasses (e.g., Lamp5, Lamp5 Lhx6, Sncg, Fig. 1E), using even small sets of ten markers in aggregate reliably recover expected trends (Fig. 1E). Notably, primate Pax6 markers are more enriched in the mouse Sncg subclass than primate Sncg markers, further supporting a misannotation across species (Fig. 1F). These findings suggest that although individual marker genes may be unreliable, aggregate subclass profiles are conserved and provide a robust basis for cross-species alignment.

We used a one-vs-best MetaNeighbor analysis to systematically map mouse interneuron clusters to primate subclasses based on transcriptomic replicability (Fig. 2A, Supp. Fig. 5). This approach identifies the best and next-best primate subclass matches for each mouse cluster and assesses how well a mouse cluster distinguishes its best primate subclass relative to its next best match, a stringent test of replicability across species. We identified robust 1-to-1 matches between mouse clusters and all nine primate subclasses, including Pax6. The homologous mouse subclass matches are represented by all 8 mouse datasets and span 18 brain regions demonstrating the strong consensus subclass alignment between primates and mice in light of significant technical and biological variability (Fig. 2B–C).

**Figure 2:**
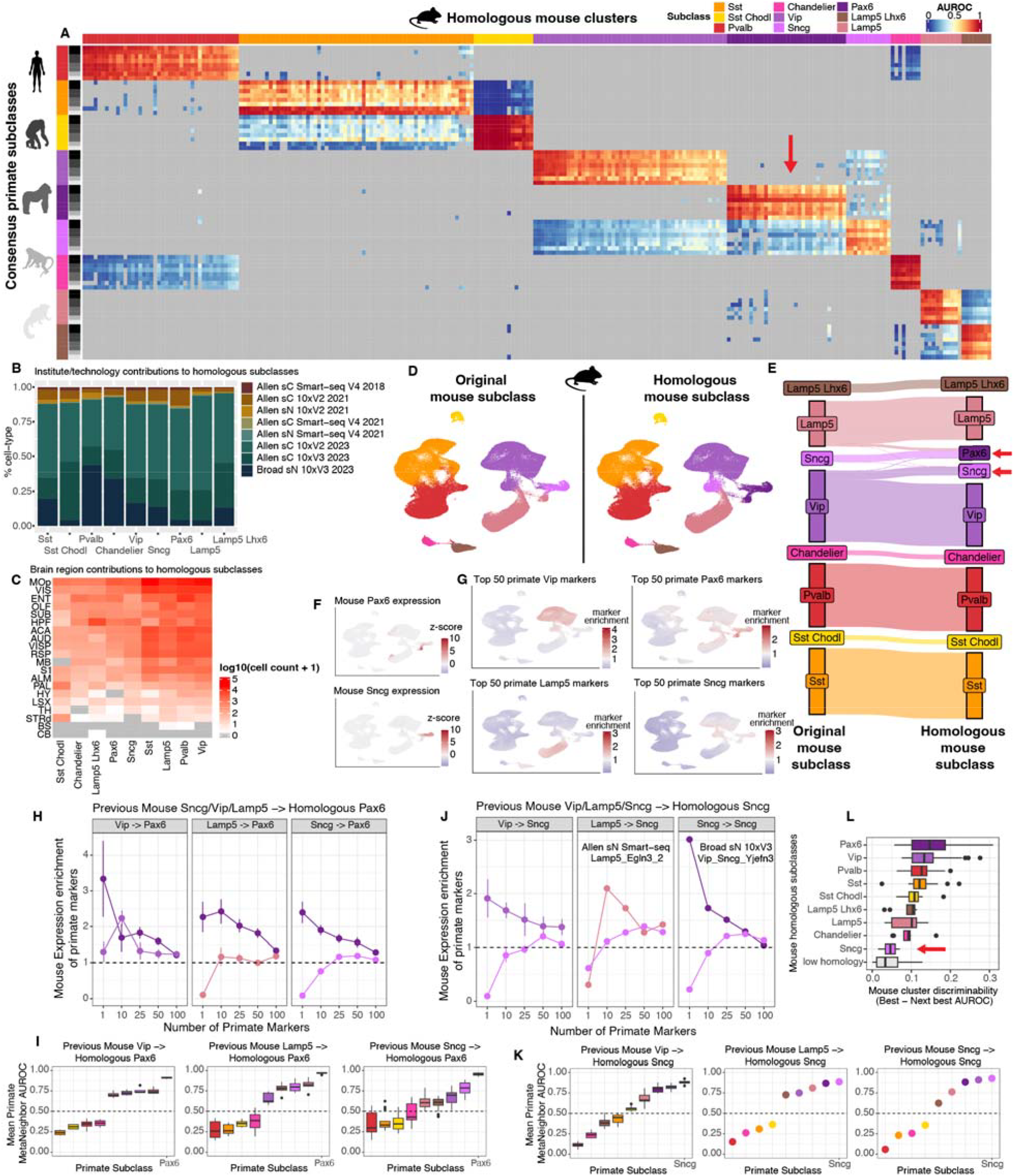
All 9 primate interneuron subclasses have homologous matches in mouse. **A:** One-vs-best MetaNeighbor scores for the replicable mouse clusters (columns) compared to the consensus primate subclasses (rows). Each mouse cluster receives an AUROC for only the top 2 subclass matches per primate dataset (16 total AUROCS for each column from the 8 total primate datasets). The red arrow is highlighting the mouse clusters aligned to the primate Pax6-subclass. Grey squares indicate mouse-cluster/primate-subclass pairs that are not in the top 2 matches per dataset comparison. **B:** Stacked barplot depicting the mouse dataset percentage composition for the 9 homologous mouse subclass annotations. **C:** Heatmap depicting the log10(cell count +1) of the 9 homologous mouse subclasses against their regional annotations. **D:** UMAPs depicting the integrated replicable mouse clusters with either their original (left) or homologous subclass annotations (right, from A). **E:** Sankey plot depicting the contributions to the homologous mouse subclass labels from the original mouse subclass labels. The red arrow is highlighting the contributions resulting in the homologous Pax6 and Sncg subclass annotations. **F:** Same UMAP as in **D**, depicting the z-scored expression of the mouse PAX6 and SNCG gene expression. **G:** Same UMAP as in **D**, depicting the marker set enrichment scores (see *Cell-type marker and MetaMarker generation* methods section) for the top 50 primate Vip, Pax6, Lamp5, and Sncg markers (Supp. Table 1) in the mouse data. **H:** Scatter-line plots comparing the marker set enrichment scores of the markers for the two associated primate subclasses of the mouse clusters that comprise the homologous Pax6 mouse subclass. For example, the left panel depicts the marker enrichment of the primate Vip and Pax6 subclass markers in the mouse clusters that switch from Vip to Pax6 subclass annotations. The points depict the mean marker enrichment score for the mouse clusters, with the lines depicting ± SD. **I:** Boxplots depicting the all-vs-all MetaNeighbor AUROC distributions of all of the primate subclasses for the mouse clusters that comprise the homologous Pax6 subclass annotations. **J:** Same as in **H**, but for the mouse clusters that comprise the homologous Sncg subclass annotations. Only a single mouse cluster represents the Lamp5 -> Sncg or the Sncg -> Sncg groups, denoted in the associated panels **K:** Same as in **I**, but for the mouse clusters that comprise the homologous Sncg subclass annotations. **L:** Boxplots depicting the distributions of differences in the best versus next best all-vs-all MetaNeighbor AUROCs comparing the mouse clusters against the primate subclasses, as a measure of discriminability between a mouse clusters best and next best primate subclass match. The grey ‘low homology’ distribution represents the scores for all of the ‘low_homology’ mouse clusters in Supp. Table 2. The red arrow highlights the mouse homologous Sncg subclass as the lowest scoring subclass, next to the low homology clusters, indicating weak replicability between the mouse and primate datasets.

To resolve the prior absence of Pax6 in mouse subclass annotations, we compared original mouse subclass labels to their primate-aligned identities (Fig. 2D–E). Nearly all previously annotated mouse Sncg clusters align instead with the primate Pax6 subclass, alongside subsets of Lamp5 and Vip clusters. Conversely, the mouse clusters most similar to the primate Sncg subclass are largely annotated as Vip. Marker expression further supported this reclassification: the majority of mouse PAX6 expression is localized to the Lamp5 cells remapped to primate Pax6, while mouse SNCG marks the Sncg-to-Pax6 clusters (Fig. 2F). This suggests the homologous primate Pax6 subclass in mice is largely composed of the previous mouse Sncg subclass along with a PAX6-expressing previous Lamp5 subset. Expanding to the top 50 primate subclass markers (Supp. Table 1) shows clear enrichment of the Vip, Lamp5, and Pax6 markers in the corresponding remapped mouse populations, while primate Sncg markers surprisingly lack strong and specific expression in the mouse data (Fig. 2G).

We explored the homologous subclass matches further with both expression enrichment (Fig. 2H) and all-vs-all MetaNeighbor analysis, which confirms Pax6 as the top primate subclass match for the associated remapped mouse clusters (Fig. 2I). In contrast, although primate Sncg remains the best overall match for its mapped mouse clusters (Fig. 2K), the weak marker enrichment (Fig. 2J) and small AUROC differences from competing subclasses (Fig. 2L) suggest poor replicability. These patterns suggest that while Pax6 is robustly conserved across species, Sncg is a weaker, potentially less reliable subclass annotation.

To further assess the replicability of the Sncg subclass, we analyzed two additional primate datasets from Yale and UT Southwestern (Fig. 3A, 3E, Supp. Fig. 4). After de novo clustering, we predicted subclass identity of individual cells using the top 50 Allen primate subclass markers (see methods). This approach yields clear 1-to-1 mappings of de novo clusters to all nine primate subclasses in both datasets (Fig. 3B, 3F), confirming that the Sncg subclass is in fact highly replicable across primate datasets.

**Figure 3:**
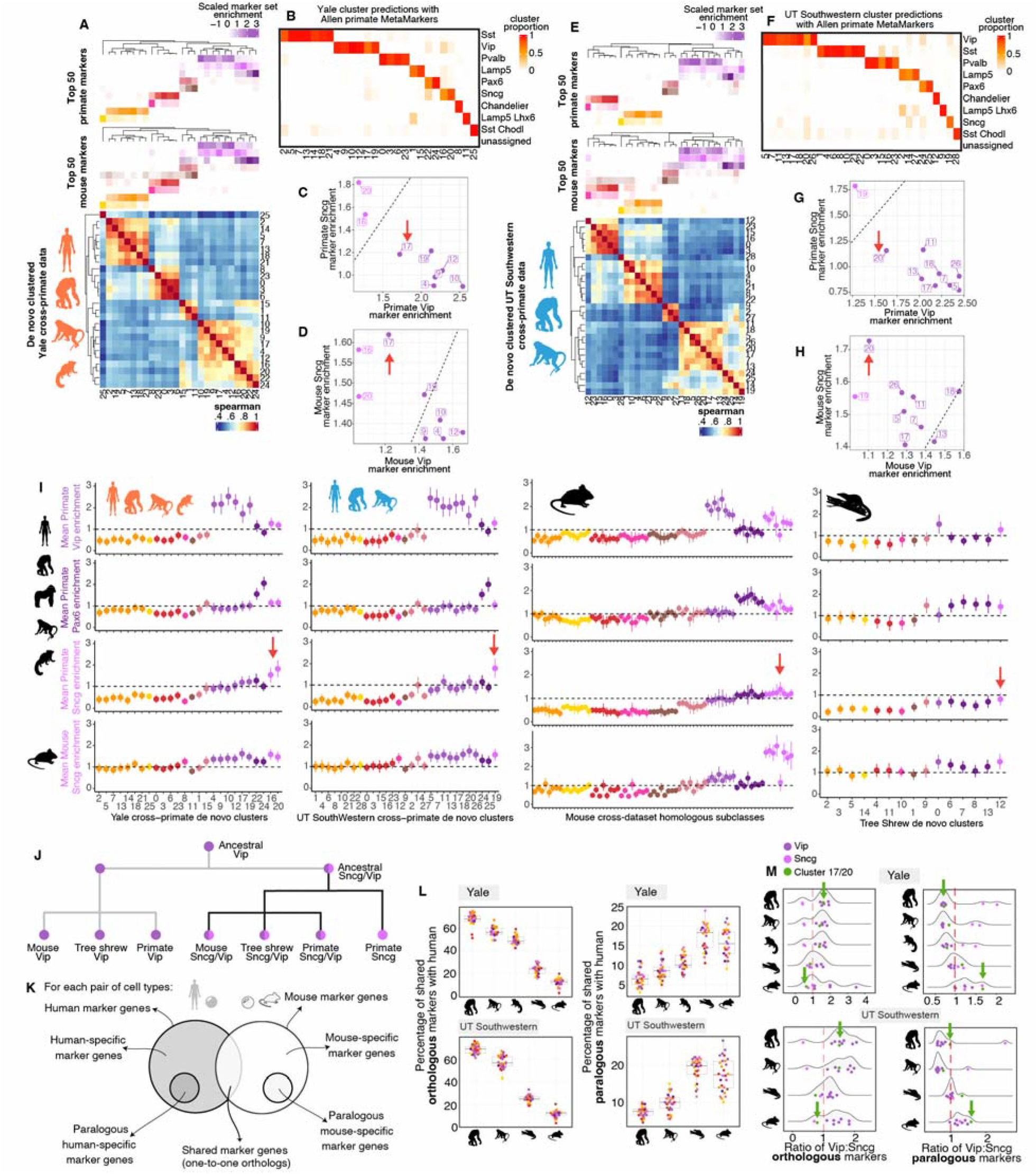
The Sncg subclass transcriptionally diverges from rodents to primates. **A:** Cluster taxonomy (see *Cell-type taxonomies* methods section) for the de novo Yale primate clusters. The heatmap depicts the spearman correlations between cluster gene expression centroids for the cluster DE genes. The column annotations depict the z-scored marker set enrichment scores for either the top 50 mouse or primate subclass markers. The color scales depict all positive z-score values. **B:** Heatmap depicting the Yale de novo cluster proportions mapped to the Allen MTG primate subclass annotations using the top 50 primate subclass MetaMarkers. **C:** Scatter plot comparing the primate Vip vs primate Sncg marker set enrichment scores for the Yale de novo clusters that map to the primate Vip and Sncg subclasses (from **B**). The red arrow highlights the Yale Vip cluster (17) with the lowest Vip marker enrichment score. **D:** Scatter plot comparing the mouse homologous Vip vs homologous Sncg marker set enrichment scores for the same Yale de novo clusters as in **C**. The red arrow highlights the same Yale Vip cluster 17 as in **C**, which has the highest enrichment for the mouse Sncg marker set. **E:** Same as in **A**, but depicting the UT Southwestern cross-primate dataset. **F:** Same as in **B**, but depicting the UT Southwestern cross-primate dataset. **G:** Same as in **C**, but depicting the UT Southwestern cross-primate dataset. **H:** Same as in **D**, but depicting the UT Southwestern cross-primate dataset. **I:** Dotplots depicting the mean (± SD) marker set enrichment for the top 50 primate Vip, Pax6, Sncg and the mouse homologous Sncg subclass markers (top to bottom) across the Yale, UT Southwestern, mouse, and tree shrew datasets (left to right). The Yale and UT Southwester de novo cluster subclass annotations are derived from panels **B** and **F**. The mouse cluster subclass annotations are derived from Figure 2A. The tree-shrew de novo cluster subclass annotations are derived from mouse and primate marker set enrichment and MetaNeighbor assessments in Supp. Fig. 7. The red arrows highlight the presence of clearly enriched primate Sncg subclass markers in the two non-Allen primate datasets, with a lack of strong primate Sncg subclass marker enrichment in the mouse and tree-shrew datasets. **J:** Model for Sncg subclass evolution. An ancestral Vip outgroup (the ancestral Sncg/Vip) is extant in mouse, tree shrew, and primates as the Vip outgroup /Sncg-like populations, with the Sncg subclass as a further diverged group within the primate lineage. **K:** Schematic outlining the orthologous (shared) and paralogous definitions for cross-species comparisons. **L:** Boxplots depicting the percentage distributions of either shared orthologous (left panels) or paralogs (right panels) between humans and non-human species for the Yale (top) and UT Southwester (bottom) datasets. **M:** Density and scatter plots depicting the ratios of either Vip:Sncg orthologous markers (left) or Vip:Sncg paralogous markers (right) for the Yale (top) and UT Southwestern (bottom) datasets. The green arrows highlight either the Yale cluster 17 or the UT Southwestern cluster 20.

We next examined how homologous mouse Sncg markers relate to primate subclasses by assessing their expression in the Yale and UT Southwestern datasets. Homologous mouse Sncg markers show intermediate enrichment in both primate Sncg and Vip clusters (Fig. 3C-D, 3G-H), with the highest enrichment found in a Vip cluster that has weak expression of primate Vip markers. This suggests that mouse Sncg markers best align with a primate Vip outgroup. Consistently, transcriptomic comparisons among mouse clusters show that homologous Sncg clusters form an outgroup relative to Vip clusters (Supp. Fig. 6).

Our comparative analyses show that the primate Sncg subclass, though highly consistent across primate datasets, has only a weak homologous match to a Vip outgroup in mouse, suggesting significant divergence between rodents and primates. To test this further, we analyzed single-cell data from tree shrew^23^—an evolutionary intermediate between rodents and primates (Supp. Fig. 7). After identifying CGE and MGE homologous cells (see methods), we assessed expression enrichment for primate Vip, Pax6, Sncg, and homologous mouse Sncg markers in tree shrew clusters (Fig. 3I). Tree shrew populations show clear enrichment for primate Vip and Pax6 subclasses, but only for the mouse, not primate, Sncg subclass – further support that the primate Sncg population represents a lineage-specific innovation.

The best homologous match for the primate Sncg subclass appears to be a Vip outgroup cluster in mouse that is also retained in tree shrew. Based on shared marker expression (Fig. 3C–D, G–H) and broader transcriptomic replicability (Supp. Fig. 4), a specific primate Vip population (Yale 17, UT Southwestern 20) aligns with this non-primate Vip outgroup /Sncg-like population. A putative explanation is that the primate Sncg subclass evolved from a Vip outgroup precursor that is retained across species - as the homologous Sncg subclass in mouse and tree shrew and as a Vip outgroup in primates (Fig. 3J).

To investigate this evolutionary trajectory, we quantified the proportion of shared orthologous versus paralogous markers across species^4^ (Fig. 3K). As expected, we observed a progressive shift from shared orthologs to shared paralogs with increasing evolutionary distance (Fig. 3L), consistent with paralog sub-functionalization contributing to cell-type divergence^4^. The human Vip outgroup clusters (Yale 17, UT Southwestern 20) switch from expressing more Vip orthologs in non-human primates to more Sncg orthologs in non-primates (Fig. 3M), reinforcing their alignment with the non-primate Sncg subclass. Concurrently, human Vip and Sncg clusters show increasing convergence on Vip paralogs - rather than Sncg paralogs - when compared to mouse (Fig. 3M) and this is specific to Vip rather than other interneuron subclasses (Supp. Fig. 7L). These data suggest the Vip and Sncg subclasses are converging transcriptionally in the transition from primates to rodents, concordant with the model that the primate Sncg subclass may be a remodeled derivative of an ancestral Vip lineage (Fig. 3J). Assessments of further diverged species from primates will be necessary to confirm.

Our analyses highlight the limitations of using small marker gene sets to define cell types across species. While reliable within species, markers can vary across evolutionary lineages^5,7^, complicating cross-species comparisons. In contrast, aggregate expression of large marker sets provides a more robust and consistent framework for aligning cell types. Using this approach, we identified a persistently misannotated CGE Pax6 subclass homolog in mouse, clarifying its absence compared to primates. We also observed notable divergence in the Sncg subclass: while consistent across primates, its best transcriptomic match in mouse and tree shrew is a Vip population, suggesting the Sncg subclass is rather a primate-specific derivation from an ancestral Vip precursor. Alongside these findings, we provide cross-mammal subclass markers (Supp. Tables 1) and revised, evolution-aware annotations for widely used datasets (Supp. Table 2). More broadly, our work shows that while within-species annotation is essential, cross-species assessments are a powerful means to address ambiguities and ground cell-types in an evolutionary framework, enabling deeper insight into both shared and lineage-specific features of neuronal diversity.

## Methods

### Data collection and processing

All data used in this work is publicly available and was sourced as provided by the original authors. We used CPM normalization for all datasets. sC: single-cell, sN: single-nucleus.

Mouse datasets:

- Allen sC Smart-seq V4 2018, Anterior Lateral motor cortex^15^
- Allen sC 10xV2 2021, Primary motor cortex^9^
- Allen sN 10xV2 2021, Primary motor cortex^9^
- Allen sC Smart-seq V4 2021, Primary motor cortex^9^
- Allen sN Smart-seq V4 2021, Primary motor cortex^9^
- Allen sC 10xV2 2023, whole brain atlas^19^
- Allen sC 10xV3 2023, whole brain atlas^19^
- Broad sN 10xV3 2023, whole brain atlas^20^

Cross-primate datasets

- Allen sN 10xV3 and Smart-seq V4 2023, Medial temporal gyrus^7^
- Yale sN 10xV3 2022, Dorsolateral prefrontal cortex^24^
- UT Southwestern sN 10xV3 2023, Posterior cingulate cortex^25^

Tree-shrew dataset

- sN 10xV3 2025, Hippocampus^23^

For the whole mouse brain atlases, the Broad sN 10xV3 2023 dataset was initially subsetted to clusters with CGE or MGE in their annotation and the Allen sC 10x 2023 datasets were initially subsetted to the CTX-MGE and CTX-CGE class annotated clusters. After the identification of replicable clusters across the 8 mouse datasets (see *MetaNeighbor* methods section), the replicable mouse clusters were integrated using the Seurat default integration approach (SelectIntegrationFeatures -> FindIntegrationAnchors -> IntegrateData) using default parameters. The integrated cell embedding was used to visualize marker gene expression in Fig. 2D, F, and G and the integrated expression data was used to construct the cluster taxonomy across datasets in Supp. Fig. 6 (see *Cell-type taxonomies* methods section).

The human, chimpanzee, gorilla, macaque, and marmoset MTG dataset was subsetted to the 31 consensus interneuron clusters previously identified across all 5 species^6,7^. This consensus subset was used to generate the primate interneuron MetaMarkers (see *Cell-type marker and MetaMarker generation* methods section) and was used in the mouse cluster vs primate subclass MetaNeighbor assessment. The primate MTG subclass annotations were sourced from Supplemental Table 6 of the associated publication^7^. The mouse datasets and the cross-primate MTG dataset were subsetted to 11,228 shared 1-to-1 orthologs across all species before any analysis; orthology information was sourced via Ensembl. All gene names across species were replaced with the human gene name equivalent.

The UT Southwestern cross-primate dataset was subsetted to the 14,649 1-to-1 orthologs identified by the previous authors. De novo clusters were computed using the author-provided cross-species integrated shared nearest neighbor graph (Supp. Fig. 4) with Seurat functions; FindNeighbors(dims = 1:25) and FindClusters(graph.name = ‘integrated_snn’) using default parameters unless otherwise noted.

The Yale cross-primate dataset was subsetted to the 28,216 1-to-1 orthologs identified by the previous authors. The data was subsetted to author-annotated ‘Inh’ cells, split by species, and then integrated using Seurat’s default approach (SelectIntegrationFeatures -> FindIntegrationAnchors -> IntegrateData). De novo clusters were computed using the cross-species integrated shared nearest neighbor graph (Supp. Fig. 4) with Seurat functions; FindNeighbors(dims = 1:25) and FindClusters(graph.name = ‘integrated_snn’) using default parameters unless otherwise noted.

The tree shrew dataset was subsetted to 9,402 1-to-1 orthologs with humans. Cells were filtered to have a minimum of 200 genes and a maximum of 6000 genes detected. Non-neuronal, Inhibitory, and Excitatory cross-primate MetaMarkers from the MTG primate consensus taxonomy^6,7^ were used to initially annotate individual cells (see *Cell-type annotation prediction with MetaMarkers* methods section). Cells were then filtered for the Inhibitory annotated cells and cells from the author-annotated ‘Infancy’ age annotation were excluded (Supp. Fig. 7). The data was then split by age (Adult and Aging) and integrated using Seurat’s default approach (SelectIntegrationFeatures -> FindIntegrationAnchors -> IntegrateData). De novo clusters were computed using the cross-age integrated shared nearest neighbor graph (Supp. Fig. 7) with Seurat functions; FindNeighbors(dims = 1:25) and FindClusters(graph.name = ‘integrated_snn’) using default parameters unless otherwise noted.

### MetaNeighbor

MetaNeighbor quantifies the strength of transcriptomic replicability for clusters across datasets using a simple supervised framework, see ref.^21^ for more details. Briefly, a cell-cell similarity graph is constructed across cells being assessed using a set of highly variable genes. We use the top 1000 highly variable genes, identified with the MetaNeighbor variableGenes() function. The cell-type annotations are hidden for a dataset (test dataset) and then predicted by their similarity to the remaining labeled datasets (training data), quantified through the AUROC metric. In the all-vs-all MetaNeighbor procedure, this produces a pairwise assessment of replicability for all the annotated cell-types across all the included datasets. This is implemented with the MetaNeighbor MetaNeighborUS(fast_version = TRUE) function with default parameters unless noted. The one-vs-best MetaNeighbor procedure first identifies the two closest matching cell types for each test dataset and computes an AUROC quantifying the replicability of the closest match relative to the next best match, as a more specific assessment of replicability between similar clusters. The one-vs-best MetaNeighbor procedure is implemented with the MetaNeighbor MetaNeighborUS(one_vs_best = TRUE, fast_version = TRUE) function.

To easily identify replicable clusters across datasets, we use a threshold on the reciprocal AUROCs between clusters. If cluster A has an AUROC of 0.95 for cluster B and cluster B has an AUROC of 0.93 for cluster A, then cluster A and B would be reciprocal best matches for any AUROC threshold <= 0.93. For the cross-mouse dataset MetaNeighbor (Supp. Fig. 3), we use a reciprocal AUROC threshold of >= 0.60 to identify replicable meta-clusters, presented in Supp. Table 3. There was a noticeable large group of replicable clusters specific to the whole brain atlas datasets (Supp. Fig. 3). A large portion of these clusters contained additional regional annotations in their cell-type labels from the Allen sC 10xV2 and 10xV3 whole brain atlas datasets, mostly with striatal tissue annotations (Supp. Fig. 3). We excluded any mouse meta-cluster that included one of these regionally annotated clusters (NDB-SI-MA-STRv, PAL-STR, STR, RHP-COA) due to these meta-clusters being exclusively comprised of clusters from the whole brain atlas datasets, including the Broad whole brain atlas (Supp. Fig. 3 and Supp. Table 3). The included meta-clusters in Supp. Table 3 comprise our replicable subset of mouse clusters used in all downstream analysis.

For several of the older mouse datasets when comparing the original subclass labels to our homologous subclass labels, we renamed specific clusters as subclasses, due to their later annotation as subclass-level cell-types. Specifically, Lamp5 Lhx6, Sst Chodl, Pvalb Vipr2, and Serpinf1 were previous clusters that we labeled as Lamp5 Lhx6, Sst Chodl, Chandelier, and Vip subclasses respectively.

We labeled the Broad whole brain atlas clusters with the Allen subclass labels (Supp. Table 4) through a majority vote of the meta-clusters in Supp. Table 3, these constitute the ‘original mouse subclass’ labels for this dataset. For the one-vs-best MetaNeighbor assessment between the replicable mouse clusters and the cross-primate MTG subclasses (Fig. 2 A), we first averaged the primate subclass AUROC scores for each mouse cluster and then called mouse clusters as high or low homologous clusters using a threshold of average primate AUROC >= 0.60 (Supp. Fig. 5). The clusters with low homology were excluded and the high homology mouse clusters were manually reordered for the plot in Fig. 2A; for each primate subclass, the mouse clusters as columns were ordered by their average primate AUROC, decreasing left to right, and the primate subclasses as rows were ordered phylogenetically.

For the Yale and Southwestern cross-primate MetaNeighbor assessments using the mouse and Allen primate data (Supp. Fig. 4) we subsetted all datasets to either the 10,637 shared 1-to-1 orthologs present across all the datasets including the Southwestern dataset or the 11,053 shared 1-to-1 orthologs shared when including the Yale dataset.

The graph plots in Supp. Fig. 4E-F and Supp. Fig 7 J-K exhibit 1-vs-best MetaNeighbor AUROCs >= 0.60 between clusters.

### Cell-type marker and MetaMarker generation

MetaMarkers is a differential expression (DE) framework that prioritizes replicable DE across datasets to rank genes as cell-type markers^10^. DE statistics are first computed for each individual dataset with the compute_markers() function; see ref^10^ for specific details. Genes are determined as cell-type DE markers for individual datasets using robust DE statistics thresholds, specifically a log2-fold change >= 4 and an FDR-adjusted p-value <= 0.05. The DE statistics are then averaged across all datasets and genes are ranked first by recurrence of DE across datasets and then by the averaged AUROC statistic to settle ties, this is performed with the make_meta_markers() function.

The primate MetaMarkers were generated from the 31 consensus interneuron clusters of the cross-primate MTG dataset using the subclass-level of annotation (Supp. Table 1). The mouse Metamarkers were generated from the 8 mouse datasets using the replicable clusters and their primate-aligned homologous subclass labels (Supp. Table 1). The cross-mammal MetaMarkers were generated using the MTG primate datasets and the 8 mouse datasets using their replicable clusters and homologous subclass labels (Supp. Table 1).

The marker enrichment score is initially computed on a per-cell basis (see ref^10^ for more details), we then average this score by cluster for the reported scores in Figs. 1E-F, Fig. 2H, J, and in Fig. 3. Let *x*_*ij*_ be the CPM-normalized expression of gene *i* in cell *j*, and *M*_*c*_ be the set of marker genes for cell type *c*. For each cell *j*, we compute a marker score as:

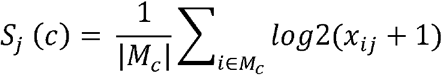

This score is the average marker gene expression of a given marker set for each individual cell, implemented by the MetaMarker score_cells() function. We then compute the marker enrichment score by first computing *S*_*j*_(*c*) for a series of cell-types *c*_1_, …, *c*_*n*_ and then compute the following for each cell-type:

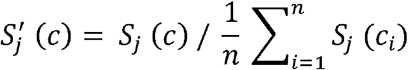

This is implemented by the MetaMarker compute_marker_enrichment() function. This normalizes marker set expression in each cell by the average expression of all marker sets, which corresponds to enrichment under the null that all marker sets are expressed equally.

### Cell-type annotation predictions with MetaMarkers

MetaMarkers uses a simple greedy approach for annotating individual cells using a collection of marker gene sets. First, marker enrichment scores are computed for each cell for a given set of cell-type markers. Then, each cell is annotated with the marker set that has the highest enrichment score; this is implemented through the MetaMarkers assign_cells() function. We use the top 50 primate subclass MetaMarkers (Supp. Table 1) to annotate the de novo clusters of the Yale and UT Southwestern cross-primate datasets (Fig. 3B, F).

### Annotating Tree shrew de novo clusters

We use the top 200 consensus primate MTG^6,7^ Inhibitory, Excitatory, and Non-neuronal MetaMarkers when initially annotating the tree shrew dataset. CGE and MGE MetaMarkers were generated using the 5 primate MTG dataset and the 8 mouse datasets using the primate-aligned homologous subclass labels (see *MetaNeighbor* methods section) and the top 100 CGE/MGE MetaMarkers were used to annotate the Inhibitory tree shrew cells as CGE or MGE (Supp. Fig. 7). The Vip, Pax6, Sncg, Lamp5, and Lamp5 Lhx6 subclasses were labeled as CGE and the Sst, Sst Chodl, Pvalb, and Chandelier subclasses were labeled as MGE. Clusters with mixed CGE-MGE proportions less than 80:20 (or 20:80) or clusters without clear enrichment of either mouse or primate subclass markers (Supp. Fig. 7) were excluded and the data was de novo clustered again for the final set of 14 clusters as in Fig. 3I. These tree-shrew clusters were annotated using a combination of mouse and primate marker enrichments and mouse vs tree shrew and primate vs tree shrew 1-vs-best MetaNeighbor assessments (Supp. Fig. 7).

### Cell-type taxonomies

Hierarchical cell-type taxonomies enable comparisons of transcriptomic similarity across groups of cell-type clusters^15–18^. Given a set of cluster annotations, the top 50 markers per cluster are first computed using the MetaMarker compute_markers() function. After filtering the expression data to the set of all top 50 markers across all clusters, the median CPM value per gene per cluster is computed as the expression centroid of each cluster. Spearman correlations are then computed across the cluster expression centroids. Hierarchical clustering using the ward.D2 algorithm is then performed on the centroid correlations, specifically using the hclust(as.dist(1-centroid_spearman), method = ‘ward.D2’) function from the R stats package, with parameters as noted. The mouse data was first filtered for only the high_homology clusters (Supp. Fig. 5, Supp. Table 3) before computing the cell-type cluster taxonomies in Supp. Fig. 6. For the integrated mouse taxonomy, we used the intersect of the top 50 markers per cluster per dataset with the 2000 highly variable genes used in the Seurat integration, for a final total of 968 genes used to construct the taxonomy.

### Computing marker overlaps between homologous cell types

Marker genes from all datasets were mapped to human gene symbols, and the list of paralogous gene pairs in humans were downloaded from Ensembl BioMart v114. For each pair of *de novo* clusters shared between human and non-human primates in Yale and UT Southwestern datasets, we compared the top 100 markers to determine the frequency of orthologous and paralogous markers expressed in related cell types (Supp. Table 5). Markers shared between a pair of clusters were defined as orthologous marker genes. Species-specific markers were identified as the genes uniquely expressed in one species after removing orthologous markers. For each species-specific gene, we retrieved its human paralogs and checked whether any were present in the orthologous marker set or in the other species’ specific markers (Fig. 3K). This analysis was performed for all cluster pairs shared between human and non-human primates. The same analysis was repeated in non-primates by comparing human clusters with *de novo* tree-shrew clusters and mouse subclasses (Supp. Table 5).

Next, we focused on Vip and Sncg clusters across species to better understand the switch in identity of Yale cluster 17 and UT Southwestern cluster 20 between primates and non-primates. For the human Vip and Sncg clusters in each dataset, we quantified the number of orthologous and paralogous marker genes shared with each Vip and Sncg cluster (in non-human primates and tree-shrew) or subclass (in mouse) in other species (Supp. Table 5). These counts were averaged by subclass and the resulting Vip:Sncg ortholog and paralog ratios were visualized to assess patterns of marker conservation and paralog substitution across species (Fig. 3M). To determine whether the increased paralog substitution observed in Yale cluster 17 and UT Southwestern cluster 20 was specific to Vip interneurons, we repeated the analysis for Sncg and non-Vip interneuron subclasses between human and mouse (Supp. Fig. 7L, Supp. Table 5).

### Software and Code Availability

All analysis was performed in R v4.4.3. Single-cell data was analyzed using either Seurat^26^ v5.0.0 objects or SingleCellExperiment^27^ v1.26.0 objects. Data visualization was performed with ggplot2 and ComplexHeatmap. All ggplot boxplot elements are the same across figure panels; the center line depicts the median, the lower and upper box limits depict the 1^st^ and 3^rd^ quartiles, and the lower and upper whiskers depict either the Q1 – 1.5*IQR or Q3 + 1.5*IQR respectively. All code used for analysis is made available at https://github.com/JonathanMWerner/mammalian_brain_consensus_taxonomy/tree/maim/code/june_2025.

## Supporting information

Supplemental Figures and legends for Supplemental Tables

## Data Availability

All data used in this work is made publicly available by the original authors, see associated citations in the *Data collection and processing* Methods section. The underlying data for all figure panels will be made available upon publication at https://github.com/JonathanMWerner/mammalian_brain_consensus_taxonomy.

## Author contributions

JMW and JG conceived the project. JMW and JG designed experiments. JMW performed analyses. HS created the Figure 1 schematics and performed the ortholog/paralog analyses. JMW wrote the draft with assistance from JG. All authors approved the final manuscript.

## Acknowledgements

JG and JMW were supported by NIH grant U24MH130968. HS was supported by NSF grant IOS-2216612. We thank all members of the Gillis lab for helpful discussions on the manuscript.

