## Supplemental Figures and legends for Supplemental Tables for "The misplaced mouse Pax6 interneuron subclass: A cross-species transcriptomic reassignment"

**Title:**

**Abstract:**

Subclass cell-types are strongly conserved across mammalian brains, making the systemic absence of a caudal ganglionic eminence (CGE) Pax6 interneuron subclass in mouse transcriptional atlases unexpected. Through cross-species transcriptomic analysis, we identify a Pax6 subclass homolog in mouse and uncover primate-specific divergence in the Sncg subclass. Our results highlight the pitfalls of relying on single marker genes and provide evolution-aware annotations and marker sets to support robust cross-mammal interneuron comparisons.

**Supplemental Table 1**

MetaMarker statistics for the primate MetaMarkers, mouse MetaMarkers, and the cross-mammal (mouse and primate datasets) MetaMarkers. Genes are initially ranked by recurrence of differential expression across datasets, then ranked by the AUROC statistic to break ties.

**Supplemental Table 2**

Original author provided cluster annotations for all mouse datasets with the homologous primate subclass annotations and whether the cluster is classified as having high or low homology (see *MetaNeighbor* methods section).

**Supplemental Table 3**

MetaNeighbor meta-cluster assignments for all mouse clusters. The excluded meta-clusters (see *MetaNeighbor* methods section) and the ‘outliers’ meta-cluster were omitted from analysis.

**Supplemental Table 4**

Mapping between the Broad cluster annotations and the original Allen subclass annotations.

**Supplemental Table 5**

Summaries of ortholog and paralog overlaps between species cluster comparisons. The ‘summary_of_shared_markers’ sheet is the ortholog and paralog percentage overlap of the top 100 markers per cluster. The ‘list_of_shared_markers’ sheet is the ortholog and paralog designation for genes. The ‘Vip_Sncg_mark_summary’ contains the overlaps from which the ratios in Fig. 3L are derived. The ‘Vip_Sncg_marker_list’ contains the ortholog and paralog designations for the Vip and Sncg clusters. The ‘human_mouse_Sncg_marker_overlap’ contains the ortholog and paralog overlaps between human and mouse cluster comparisons as in Supp. Fig.L.

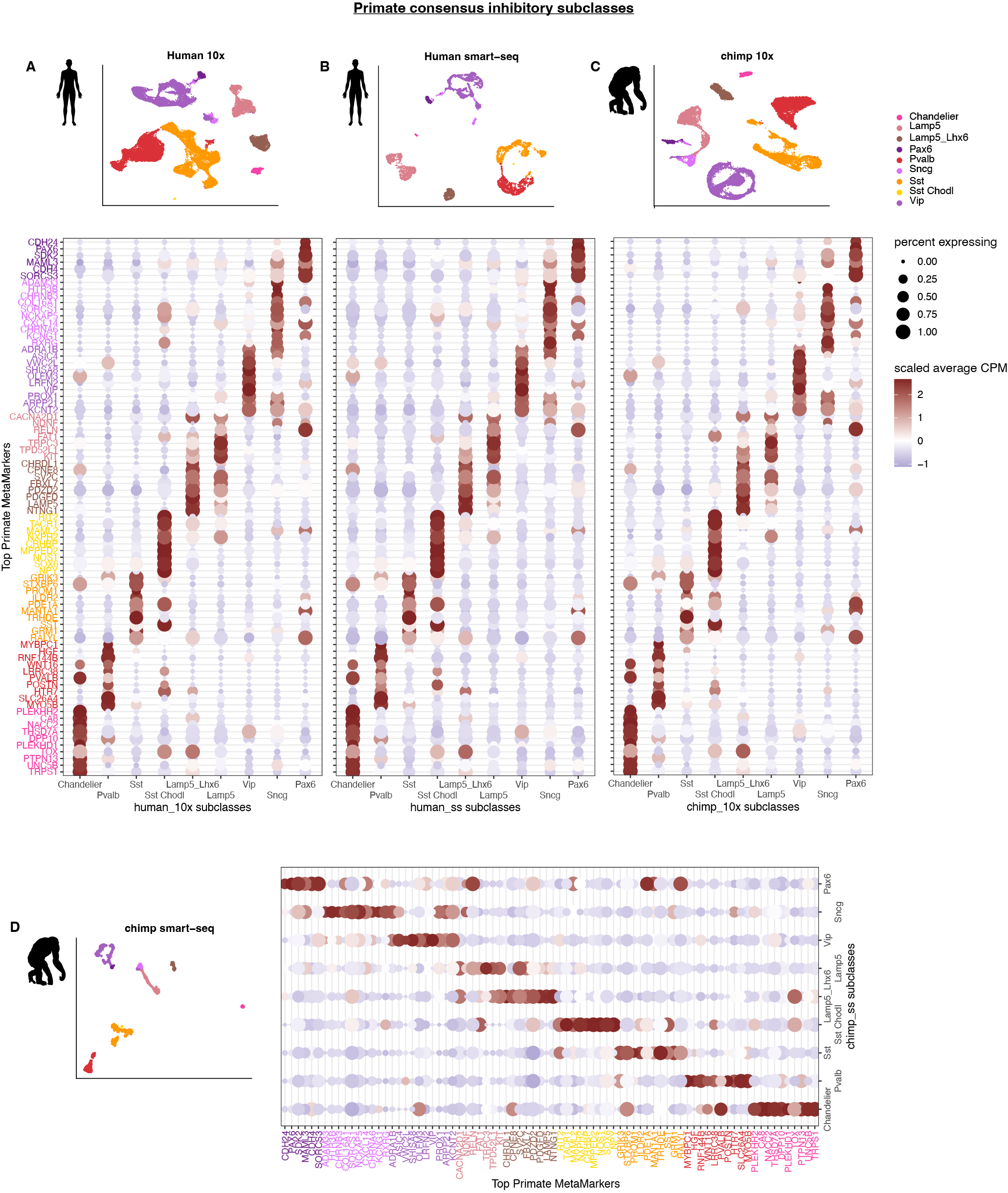

**Supplemental Figure 1: Consensus cross-primate interneuron subclass marker gene expression for human and chimpanzee datasets**

**A:** UMAP depicting the cross-primate consensus subclass annotations for the human MTG 10x dataset. Dotplot depicting the z-scored CPM values (color) and the percentage of cells expressing (CPM > 0, dot size) the top 10 primate subclass MetaMarkers (Supp. Table 1) across the subclass annotations for the human MTG 10X dataset. Duplicate MetaMarkers across the top 10 are colored for visualization purposes.
**B:** Same as in **A**, but for the human Smart-seq MTG dataset.
**C:** Same as in **A**, but for the chimpanzee 10x MTG dataset.
**D:** Same as in **A**, but for the chimpanzee Smart-seq MTG dataset.

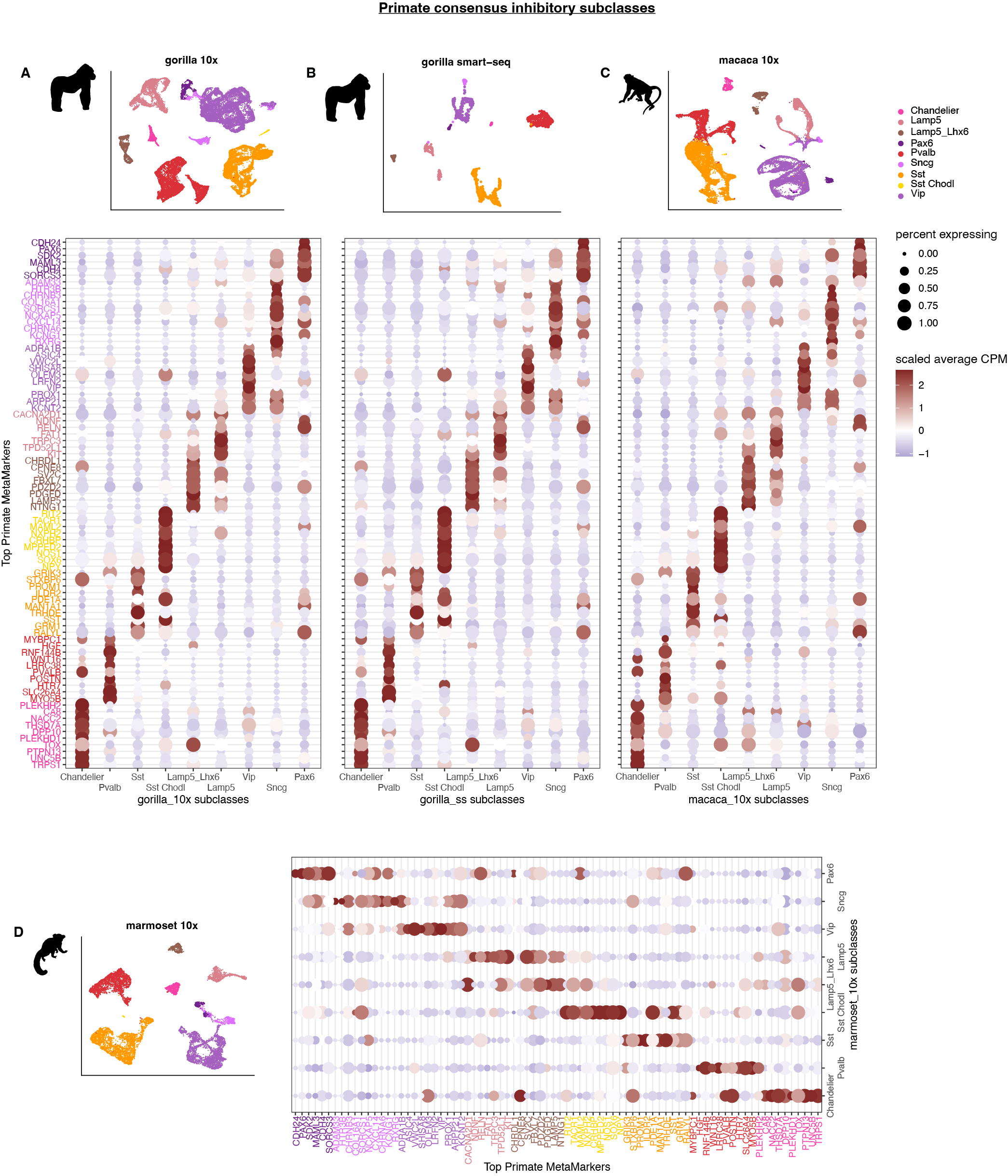

**Supplemental Figure 2: Consensus cross-primate interneuron subclass marker gene expression for gorilla, macaque, and marmoset datasets**

**A:** UMAP depicting the cross-primate consensus subclass annotations for the gorilla MTG 10x dataset. Dotplot depicting the z-scored CPM values (color) and the percentage of cells expressing (CPM > 0, dot size) the top 10 primate subclass MetaMarkers (Supp. Table 1) across the subclass annotations for the gorilla MTG 10X dataset. Duplicate MetaMarkers across the top 10 are colored for visualization purposes.
**B:** Same as in **A**, but for the gorilla Smart-seq MTG dataset.
**C:** Same as in **A**, but for the macaque 10x MTG dataset.
**D:** Same as in **A**, but for the marmoset 10x MTG dataset.

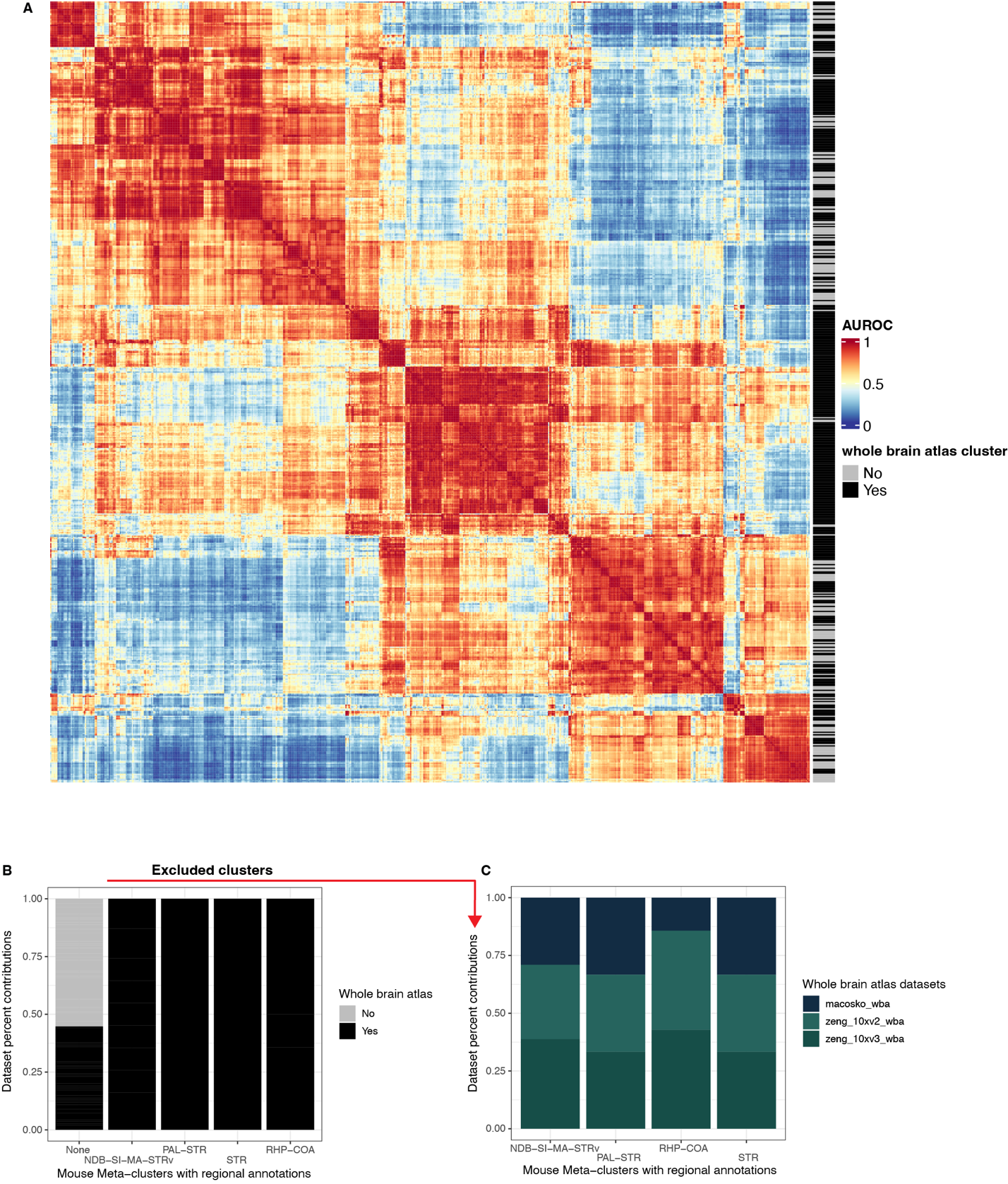

**Supplemental Figure 3: Replicability of mouse clusters**

**A:** Heatmap depicting the all-vs-all MetaNeighbor AUROC scores for all clusters across the 8 mouse datasets. The row annotation denotes whether that cluster belongs to one of the whole brain mouse datasets. Note the large collection of replicable clusters nearly exclusively from the whole brain atlas datasets.
**B:** Stacked barplots depicting the dataset contributions (whole brain atlas or not) of mouse meta-clusters (see *MetaNeighbor* methods section) that include clusters with either NDB-SI-MA-STRv, PAL-STR, STR, RHP-COA or none of these regional annotations. The meta-clusters that include those regional annotations are exclusively composed of clusters from the whole-brain datasets.
**C:** Stacked barplots depicting the dataset contributions (dataset labels) of those meta-clusters that include the regional annotations. All 3 whole brain datasets contribute to these meta-clusters, and these clusters are excluded from the mouse datasets for all downstream analysis.

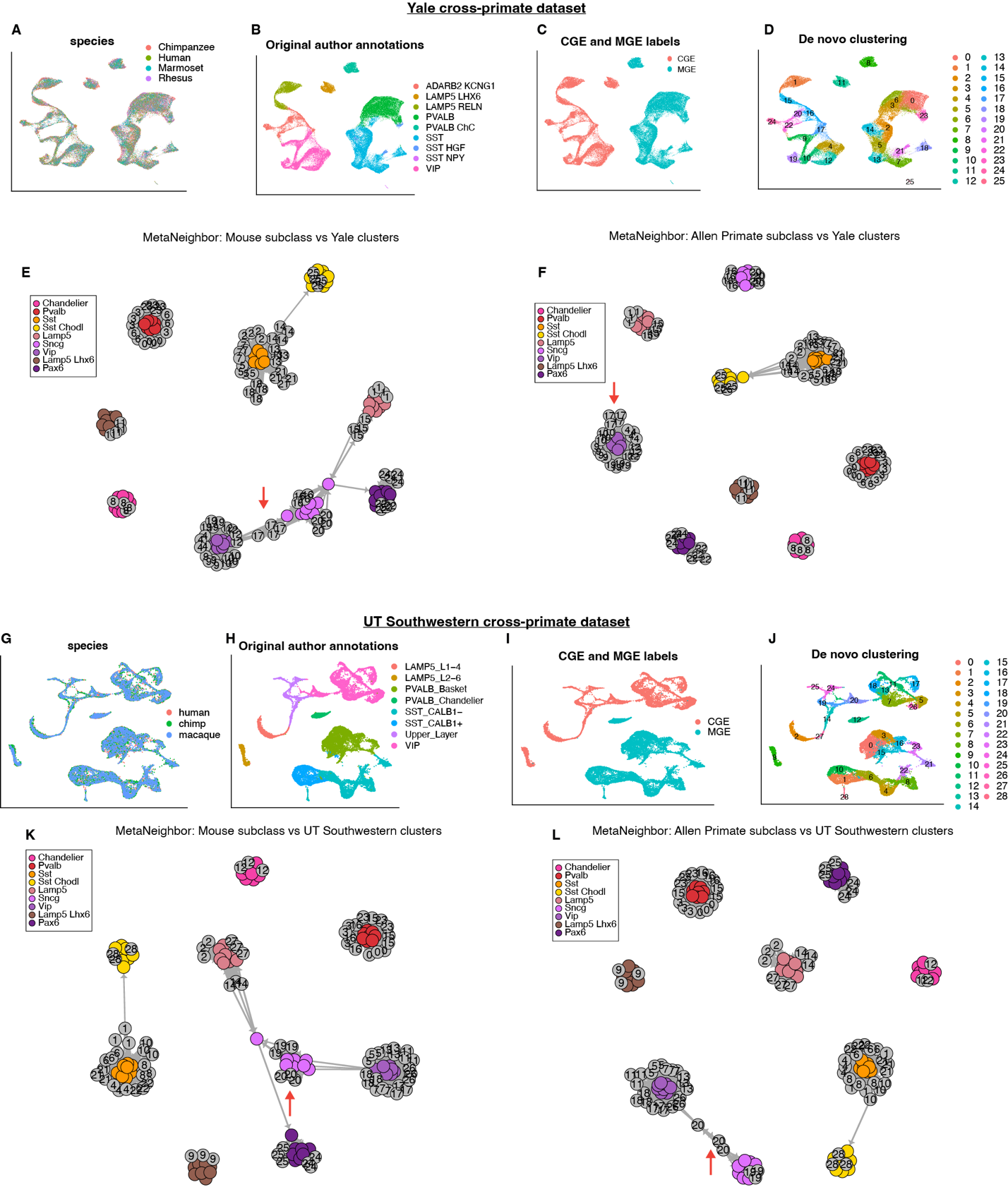

**Supplemental Figure 4: Yale and UT Southwestern cross-primate dataset de novo clustering and subclass MetaNeighbor assessments**

**A:** UMAP depicting the integrated cross-species data for the Yale dataset, colored by species.
**B:** Same UMAP as in **A**, colored by the authors original cell-type annotations.
**C:** Same UMAP as in **A**, colored by the CGE and MGE classification of the author cell-type annotations.
**D:** Same UMAP as in **A**, depicting the de novo clusters of the Yale dataset.
**E:** Graph plot depicting the one-vs-best MetaNeighbor scores between the mouse homologous subclasses and the Yale de novo clusters. Edges denote AUROCs >= 0.60, edge length corresponds inversely to AUROCs (high AUROCs -> small edges). The red arrow highlights cluster 17, which has intermediate primate Vip/mouse Sncg characteristics (from Figure 3).
**F:** Same as in panel **E**, but depicting the MetaNeighbor scores between the Allen MTG primate subclasses and the Yale de novo clusters.
**G:** UMAP depicting the integrated cross-species data for the UT Southwestern dataset, colored by species.
**H:** Same UMAP as in **G**, colored by the authors original cell-type annotations.
**I:** Same UMAP as in **G**, colored by the CGE and MGE classification of the author cell-type annotations.
**J:** Same UMAP as in **G**, depicting the de novo clusters of the UT Southwestern dataset.
**K:** Graph plot depicting the one-vs-best MetaNeighbor scores between the mouse homologous subclasses and the UT Southwestern de novo clusters. Edges denote AUROCs >= 0.60, edge length corresponds inversely to AUROCs (high AUROCs -> small edges). The red arrow highlights cluster 20, which has intermediate primate Vip/mouse Sncg characteristics (from Figure 3).
**L:** Same as in panel **K**, but depicting the MetaNeighbor scores between the Allen MTG primate subclasses and the UT Southwestern de novo clusters.

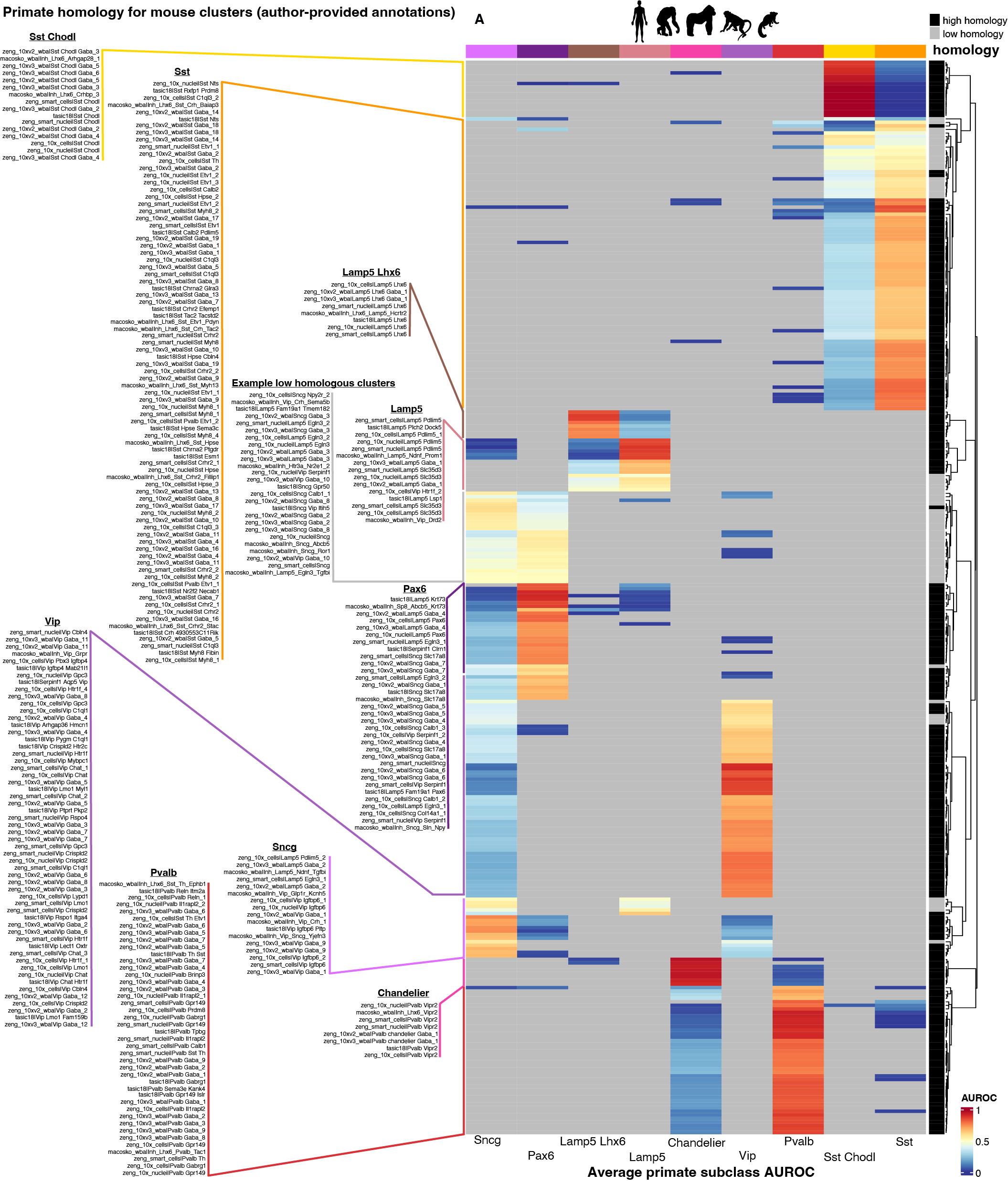

**Supplemental Figure 5: Average primate subclass MetaNeighbor AUROCs for all replicable mouse clusters**

**A:** Heatmap depicting the average primate subclass one-vs-best MetaNeighbor AUROC (columns) for all of the replicable mouse clusters (rows). The row annotation denotes the mouse clusters with either high (an AUROC >= 0.60) or low (an AUROC < 0.60) homology. The author annotations for all of the replicable mouse clusters are to the left of the heatmap, denoted by their mapped homologous primate subclass annotation. Full annotations are present in Supp. Table 2.

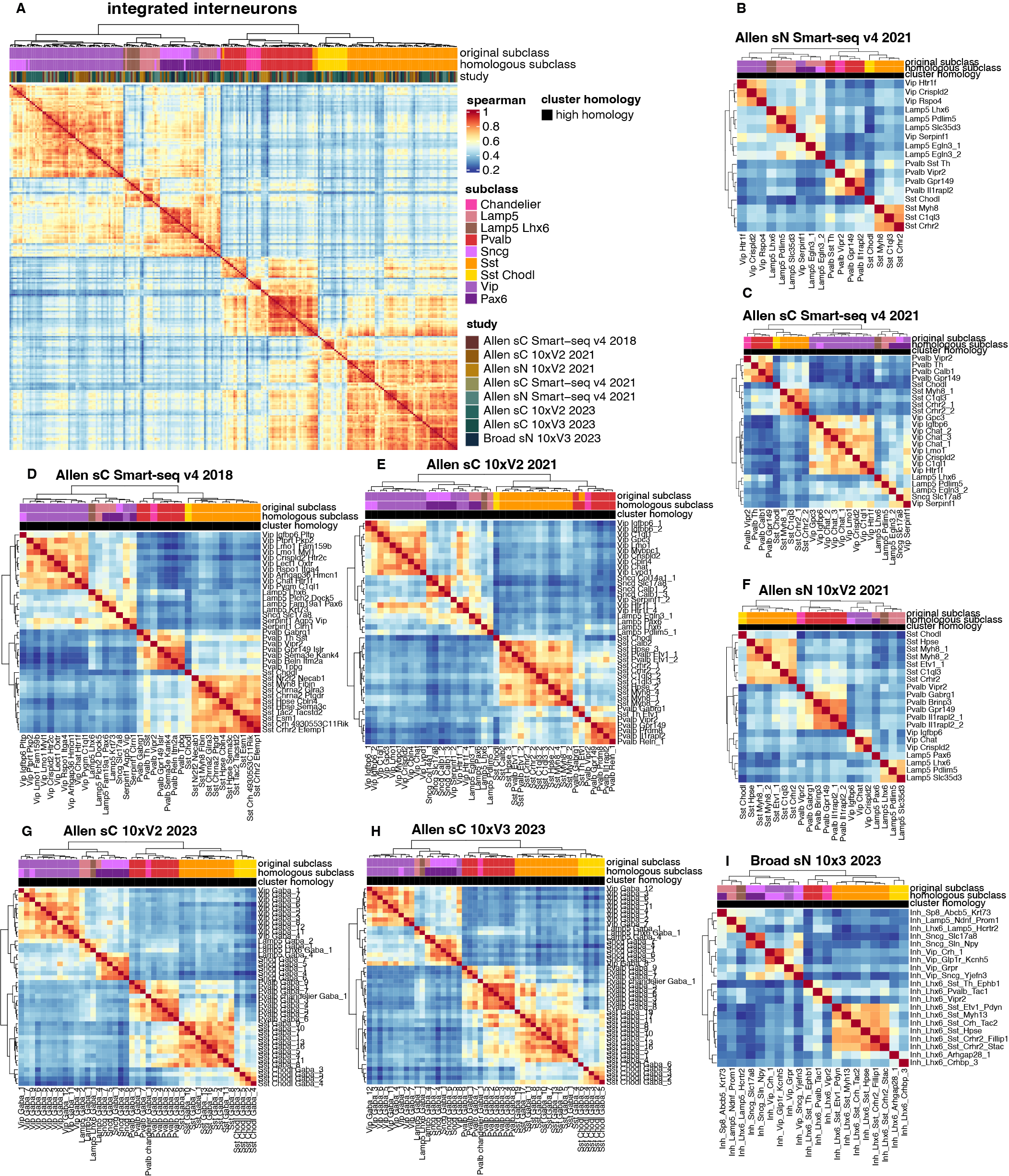

**Supplemental Figure 6: Mouse cell-type taxonomies and original vs homologous subclass annotations**

**A:** Cell-type taxonomy (see *Cell-type taxonomies* methods section) of the integrated replicable mouse clusters with high primate homology.
**B:** Cell-type taxonomy of the Allen sN Smart-seq v4 2021 mouse clusters with high primate homology, using original author annotations.
**C:** Cell-type taxonomy of the Allen sC Smart-seq v4 2021 mouse clusters with high primate homology, using original author annotations.
**D:** Cell-type taxonomy of the Allen sC Smart-seq v4 2018 mouse clusters with high primate homology, using original author annotations.
**E:** Cell-type taxonomy of the Allen sC 10x V2 2021 mouse clusters with high primate homology, using original author annotations.
**F:** Cell-type taxonomy of the Allen sN 10x V2 2021 mouse clusters with high primate homology, using original author annotations.
**G:** Cell-type taxonomy of the Allen sC 10x V2 2023 mouse clusters with high primate homology, using original author annotations.
**H:** Cell-type taxonomy of the Allen sC 10x V3 2023 mouse clusters with high primate homology, using original author annotations.
**I:** Cell-type taxonomy of the Broad sN 10x V3 2023 mouse clusters with high primate homology, using original author annotations.

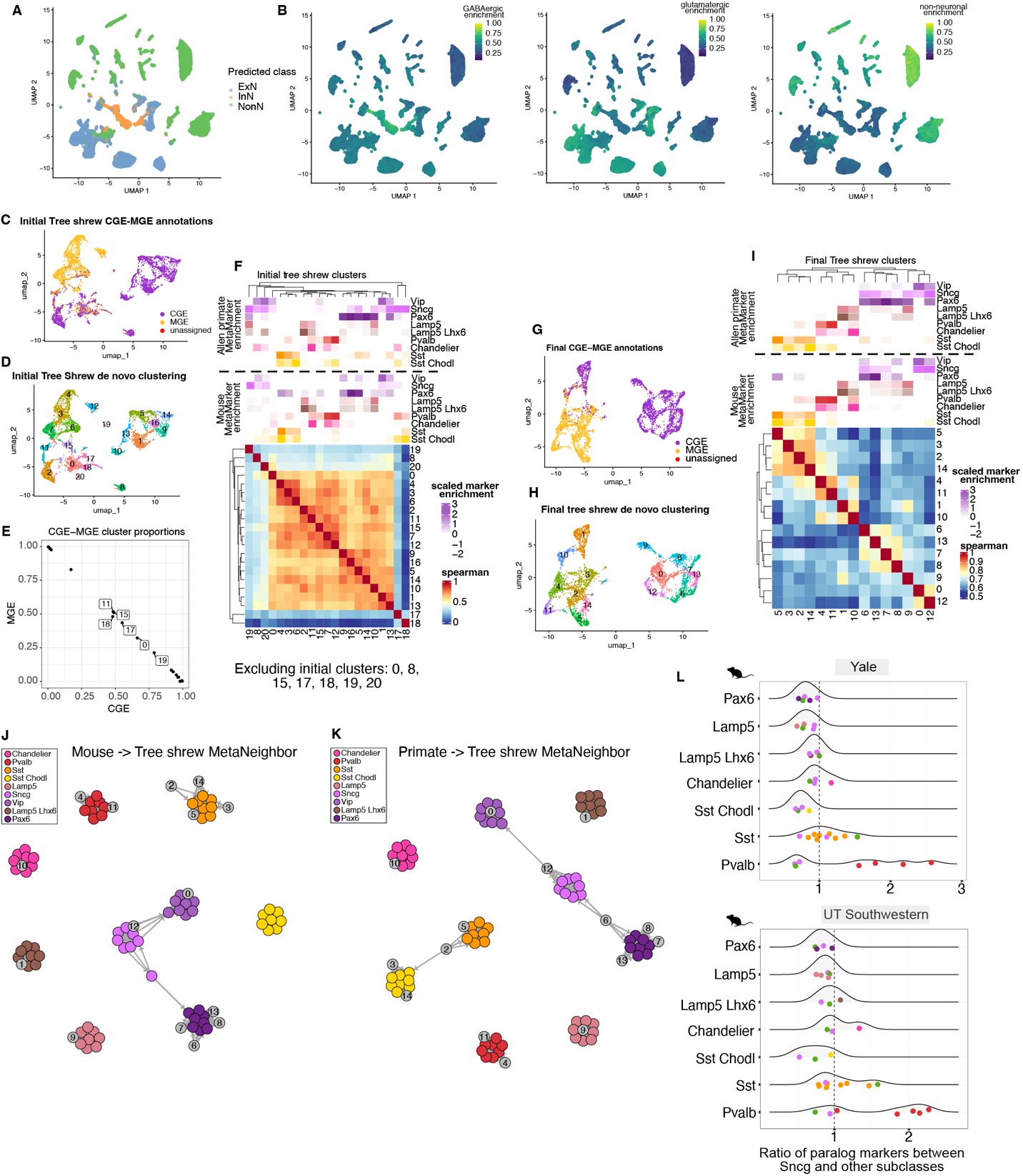

**Supplemental Figure 7: Tree-shrew processing and mouse paralogous marker cell-type ratios**

**A:** UMAP depicting the full tree-shrew dataset with the initial Excitatory, Inhibitory, and Non-neuronal annotations (see the *Annotating Tree shrew de novo clusters* methods section)
**B:** Same UMAP as in **A**, but depicting either the Excitatory, Inhibitory, or Non-neuronal MetaMarker enrichment scores, max-normalized.
**C:** UMAP depicting the Inhibitory cells annotated as CGE or MGE (see the *Annotating Tree shrew de novo clusters* methods section).
**D:** Same UMAP as in **C**, but depicting the de novo clusters.
**E:** Scatter plot comparing the CGE and MGE cluster proportions for the de novo clusters. Clusters with less than 80% CGE or MGE composition are labeled. All of these CGE-MGE intermediate clusters were excluded, with the exception of cluster 11 due to its clear high enrichment of Chandelier marker expression, in panel **F**.
**F:** Cell-type taxonomy of the tree shrew de novo clusters with the associated mouse or primate subclass MetaMarker expression enrichment scores, as in Figure 3 A and E. Clusters 8 and 20 were additionally excluded due to their poor transcriptomic similarity to other clusters and a lack of clear enrichment for any of the subclass labels.
**G:** UMAP depicting the final CGE and MGE annotations after the cluster filtering
**H:** Same UMAP as in **G**, but depicting the final set of de novo clusters.
**I:** Cell-type taxonomy of the final tree-shrew de novo clusters with the associated mouse or primate subclass MetaMarker expression enrichment scores.
**J:** Graph plot depicting the one-vs-best MetaNeighbor scores between the mouse homologous subclasses and the tree-shrew de novo clusters. Edges denote AUROCs >= 0.60, edge length corresponds inversely to AUROCs (high AUROCs -> small edges).
**K:** Same as in **G**, but depicting the Allen MTG primate subclasses and the tree-shrew de novo clusters.
**L:** Density and scatter plots depicting the ratios of paralogous markers for human vs mouse subclass comparisons. Same plot as in Figure 3L for the mouse, but with all the other subclasses other than Vip against the Sncg subclass.
